# Asymmetric neuromodulation in the respiratory network contributes to rhythm and pattern generation

**DOI:** 10.1101/2024.11.11.623076

**Authors:** Rishi R. Dhingra, Peter M. MacFarlane, Peter J. Thomas, Julian F.R. Paton, Mathias Dutschmann

**Affiliations:** Division of Pulmonary, Critical Care and Sleep, Department of Medicine, Case Western Reserve University, Cleveland, OH, USA; The Florey Department of Neuroscience & Mental Health, University of Melbourne, Parkville, Victoria, Australia; Department of Pediatrics, Rainbow Babies & Children’s Hospital, Case Western Reserve University, Cleveland, OH, USA; Department of Mathematics, Applied Mathematics and Statistics, Case Western Reserve University, Cleveland, OH, USA; Manaaki Manawa – The Centre for Heart Research, Department of Physiology, University of Auckland, Auckland, New Zealand

**Author notes:** These authors share senior authorship.

**Keywords:** breathing, opioid-induced respiratory depression, multi-electrode array, Hopfield network, rhythm generation, central pattern generator

## Abstract

Like other brain circuits, the brainstem respiratory network is continually modulated by neurotransmitters that activate slow metabotropic receptors. In many cases, activation of these receptors only subtly modulates the respiratory motor pattern. However, activation of some receptor types evokes the arrest of the respiratory motor pattern as can occur following the activation of µ-opioid receptors. We propose that the varied effects of neuromodulation on the respiratory motor pattern depend on the pattern of neuromodulator receptor expression and their influence on the excitability of their post-synaptic targets. Because a comprehensive characterization of these cellular properties across the respiratory network remains challenging, we test our hypothesis by combining computational modelling with ensemble electrophysiologic recording in the pre-Bötzinger complex (pre-BötC) using high-density multi-electrode arrays (MEA). Our computational model encapsulates the hypothesis that neuromodulatory transmission is organized asymmetrically across the respiratory network to promote rhythm and pattern generation. To test this hypothesis, we increased the strength of neuromodulatory connections in the model and used selective agonists *in situ* while monitoring pre-BötC ensemble activities. The model predictions of increasing slow inhibition were consistent with experiments examining the effect of systemic administration of the 5HT1aR agonist 8-OH-DPAT. Similarly, the predicted effects of increasing slow excitation in the model were experimentally confirmed in pre-BötC ensemble activities before and after systemic administration of the µ-opioid receptor agonist fentanyl. We conclude that asymmetric neuromodulation can contribute to respiratory rhythm and pattern generation and accounts for its varied effects on breathing.

## Introduction

Neuromodulation is essential for adaptive function in brain circuits (1–4). Neuromodulatory transmitters act via metabotropic receptors coupled to intracellular signaling cascades to slowly modify synaptic and membrane properties thereby altering circuit computations mediated by fast synaptic neurotransmission (5). For example, phasically active midbrain dopaminergic neurons encode a reward prediction error signal that modulates excitability, and hence, activity-dependent plasticity in their target populations (1).

Given the coordination of breathing with other orofacial behaviors including swallowing, vocalization and autonomic regulation, it is not surprising that many neuromodulators influence the breathing motor pattern through their actions on the brainstem respiratory network including, but not limited to serotonin, dopamine, acetylcholine, opioids, histamine, substance P and somatostatin (6–9). Neurons which express pre- and post-synaptic receptors for neuromodulatory neurotransmission are highly distributed across the respiratory network. However, the effects of neuromodulation on breathing are commonly investigated at either a coarse scale by examining their effects on the frequency and amplitude of respiratory motor nerve activities after systemic drug application or at a finer scale via drug micro-injection within a particular compartment of the respiratory network. Consequently, the mechanisms of respiratory neuromodulation identified by these experimental approaches have highlighted the role of neuromodulation within single network compartments, especially the pre-Bötzinger complex (6,9), as the primary targets of neuromodulators. However, these studies do not consider the pattern of neuromodulatory neurotransmission across the entire network, which was a major aim of the present study.

The conundrum of neuromodulation is highlighted by research concerned with opioid-induced respiratory depression (ORID) evoked by overdose of opioid-based analgesics or drugs of abuse that predominantly bind to the µ-opioid receptor (µ-OR) (10–12). Mechanistically, µ-OR agonists have been shown to act on the medullary pre-BötC (13–15), ventral respiratory group (16–20), and pontine parabrachial and Kölliker-Fuse nuclei (21–26). In addition to these functionally identified areas, a recent anatomical study of *Oprm1* expression in the respiratory network has identified µ-OR^+^ neurons in the nucleus tractus solitarii, Bötzinger complex, intermediate reticular nucleus/post-inspiratory complex, parafacial area, locus coeruleus and raphé nuclei (27) suggesting that opioids may act simultaneously at many diverse sites across the brainstem respiratory network. Despite the widespread expression of µ-ORs, several studies have proposed that OIRD depends solely on the activation of µ-ORs in the pre-BötC to suppress inspiratory rhythm generation (11,13,28). Other studies have taken a more holistic view acknowledging the role of a distributed network mechanism for OIRD, but highlight the Kölliker-Fuse nuclei as a primary therapeutic target for OIRD (10,23,24,26).

This on-going debate has motivated the need to develop an understanding of the network mechanism of OIRD (12), and of respiratory network neuromodulation, in general. However, understanding the network mechanisms for neuromodulation in a distributed brain circuit would require defining not only the complete connectome of the circuit, but also the pattern of neuromodulatory co-transmitters and receptors expression across that connectome (2,5). Here, to overcome this challenge, we combine computational modelling with ensemble electrophysiology to test the hypothesis that neuromodulatory systems in the respiratory network are organized to contribute to the maintenance of the breathing rhythm and pattern. To encapsulate this hypothesis, we follow the approach of Kleinfeld and Sompolinsky (29) who developed a pair of Hebbian learning rules for the fast- and slow-synapses of a Hopfield network that produce the periodic sequential activities observed in central pattern generating networks. By training such a model to produce the respiratory firing patterns observed in the pre-Bötzinger complex (pre-BötC), an essential node of the respiratory network that expresses a representative set of firing patterns associated with all three phases of the breathing pattern under intact network conditions *in vivo* (30–32) and *in situ* (33), we develop a model of the respiratory network in which the asymmetric pattern of slow-/neuromodulatory-connectivity drives the respiratory rhythm and pattern. To test our hypothesis, we compare the *in silico* predictions of increasing the strength of subsets of neuromodulatory connections based on the net effect on their post-synaptic targets with electrophysiologic experiments *in situ* in which we used a high-density multi-electrode array to monitor the ensemble activities of pre-BötC neurons before and after systemic administration of either the G_i/o_-coupled µ-OR agonist fentanyl or the G_i/o_-coupled 5HT1A receptor agonist 8-OH DPAT. In either case, we observed qualitatively similar responses of network activity to the perturbations in both simulations and experiments. Interestingly, the model also predicted the existence of a population code in which network activity is maximal at the transitions between the three phases of the breathing pattern, which we also observed experimentally in the ensemble activity of the pre-BötC. Taken together, we propose that neuromodulatory systems of the respiratory network are organized asymmetrically to contribute to the maintenance of the breathing rhythm and pattern. Furthermore, we conclude that activation of µ-ORs disrupts a network mechanism of respiratory rhythm and pattern generation.

## Materials and Methods

Experimental protocols were approved by and conducted with strict adherence to the guidelines established by the Animal Ethics Committee of the Florey Department of Neuroscience & Mental Health, University of Melbourne, Melbourne, Australia (AEC No.: 17-075-FINMH). For breeding, adult male and female Sprague– Dawley rats (Animal Resources Centre, Canning Vale, Australia) and their offspring were housed under a 14:10 light/dark cycle with ad libitum access to standard laboratory chow and water.

### In situ arterially-perfused brainstem preparation

Experiments were performed in juvenile (17-26 days post-natal) Sprague-Dawley rats of either sex using the *in situ* arterially-perfused brainstem preparation as described previously (34,35). Briefly, rats were anesthetized by inhalation of isoflurane (2-5%) until they reached a surgical plane of anesthesia. Next, rats were transected sub-diaphragmatically and immediately transferred to an ice-cold bath of artificial cerebrospinal fluid (aCSF; in mM: [NaCl] 125, [KCl] 3, [KH2PO4] 1.25, [MgSO4] 1.25, [NaHCO3] 24, [CaCl2] 2.5 and D-glucose 10) for decerebration. Next, the heart and lungs were removed. The phrenic nerve was isolated for recording, and the descending aorta was isolated for cannulation. Next, the cerebellum was removed. Finally, the vagus and hypoglossal nerves were isolated for recording.

The preparation was then transferred to a recording chamber. The aorta was quickly cannulated with a double-lumen catheter. The preparation was then re-perfused with carbogenated (95%/5% pO2/pCO2), heated (31°C) aCSF (200 mL) using a peristaltic pump (Watson-Marlow).

Phrenic, vagal and hypoglossal nerves were mounted on suction electrodes to record the fictive respiratory motor pattern. Motor nerve potentials were amplified (400×), filtered (1-7500 Hz), digitized (30 kHz) via a 16-channel differential headstage (Intan RHD2216), and stored on an acquisition computer using an Open-Ephys acquisition system (Rev. 2, (Siegle et al., 2017)). Within minutes, apneustic respiratory contractions resumed.

The perfusion flow rate was adjusted to fine tune the preparation to generate a stable rhythm with augmenting inspiratory phrenic discharge and bi-phasic inspiratory and post-inspiratory activity on the vagus nerve. Finally, a single bolus of vecuronium bromide (0.3 mL, 0.1 mg/mL w/v vecuronium bromide: saline) was delivered to the perfusate to paralyse the preparation to avoid movement artifacts.

### Ensemble recording of pre-Bötzinger complex

In one series of experiments (n=11), we measured single unit activities across ensembles of pre-BötC neurons using a 4-shank, 64-channel high-density silicon MEA (Neuronexus, A4×16) while simultaneously recording the three-phase respiratory motor pattern on phrenic, vagal and hypoglossal nerves. The MEA electrode sites spanned 345 µm in the dorso-ventral axis, and 600 µm in the rostro-caudal axis.

Using a micro-manipulator (Narishige MMN-33), we slowly inserted the MEA into the brainstem until we observed an ensemble of neuronal activities with respiratory-related firing patterns. The coordinates of the recording sites were measured from the caudal-most shank relative to those of calamus scriptoriius and were: 1.6-2.3 mm rostral to calamus scriptorius, 1.4-1.8 mm lateral to the midline and 1.6-2.2 mm below the brainstem surface. Once positioned within the pre-BötC, we recorded the spontaneous activity of the pre-BötC ensemble for 10 min. Neuronal activities from the MEA were amplified (400×), filtered (0.001-7.5 kHz) and digitized via a 64-channel mono-polar headstage (Intan RHD2164) and stored on an acquisition computer using an Open-Ephys acquisition system.

In a subset of these experiments, to enable mapping the location of pre-BötC neurons to the 7T MRI Waxholm atlas of the Sprague-Dawley rat brain (36) by determining the rigid transformation necessary to register the positioner coordinate-system with those of the Waxholm atlas, we measured the coordinates of 5 surface landmarks that were easily identifiable both on the brainstem surface of the preparation and within the atlas (Suppl. Fig. 1, Fig. 1A & F).

**Figure 1:**
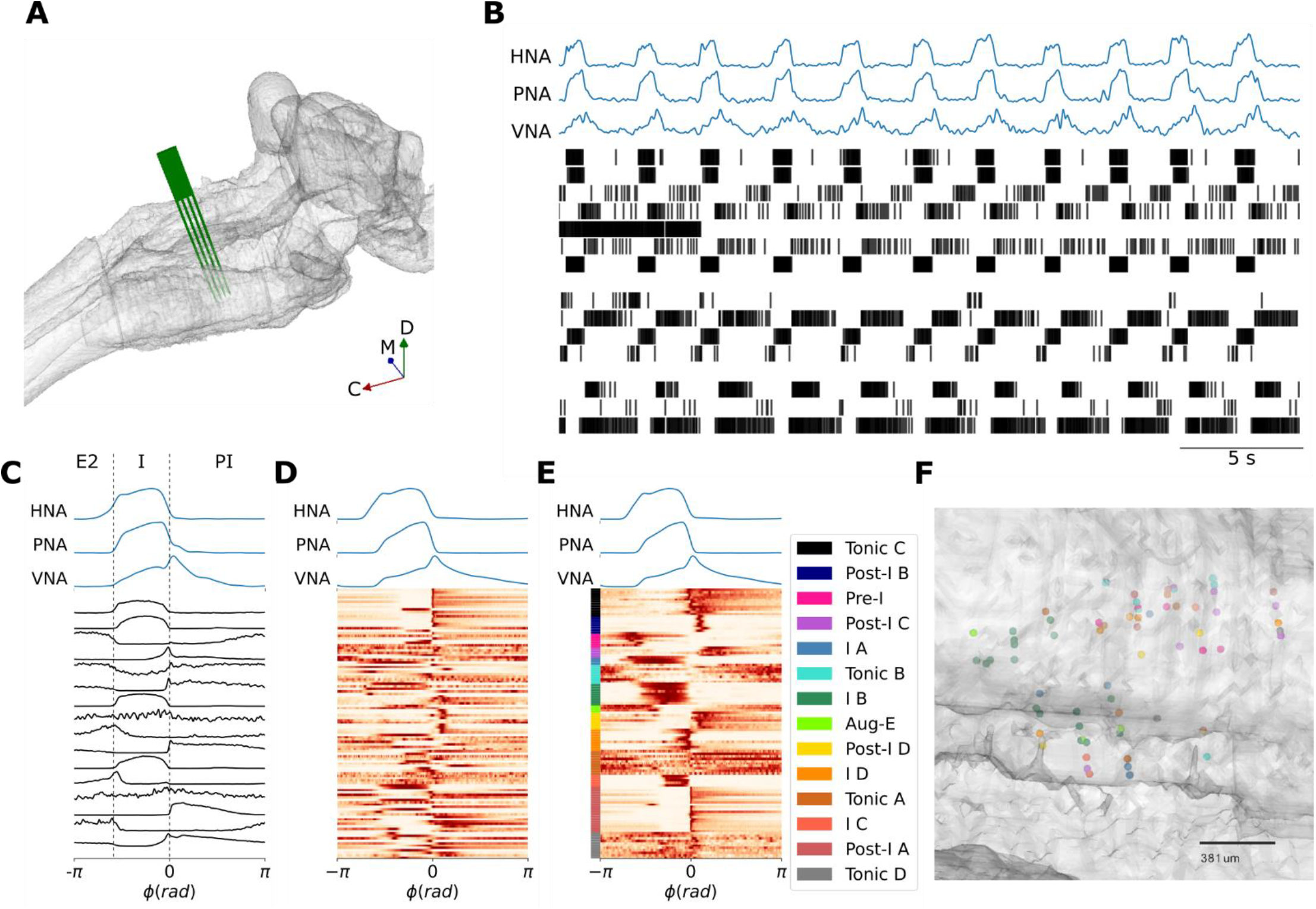
Identifying the pre-BötC neuronal firing patterns that underlie the three-phase respiratory motor pattern. **A** Reconstruction of an MEA positioned in the pre-BötC *in situ* in a representative experiment. **B** A representative recording of the three-phase respiratory motor pattern on hypoglossal (HNA), phrenic (PNA) and vagal (VNA) nerves in concert with an ensemble of pre-BötC neurons. **C** Cycle-triggered histograms of the pre-BötC neurons shown in (**A)** include neurons that spike in late-expiration (E2), inspiration (I) and post-inspiration (PI). **D** Cycle-triggered histograms of all recorded pre-BötC neurons before clustering. **E** K-means clustering identifies 14 classes of pre-BötC firing patterns that underlie the three-phase respiratory motor pattern. **F** Reconstructed locations of a subset of neurons suggest that pre-BötC neuronal types are spatially mixed.

### Pharmacologic experiments

In subsequent experiments, to assess the effects of increasing neuromodulatory tone, after positioning the MEA within the pre-BötC and recording the stationary baseline pattern of pre-BötC ensemble activity, we administered either the 5HT1aR agonist 8-OH DPAT (1 µM, n=4) or the µ-opioid receptor agonist fentanyl (15 nM, n=8) to the perfusate and recorded the ensemble activity of the pre-BötC for an additional 10 min once the preparation expressed a new stationary breathing pattern (≤ 5 min).

### Data Analysis

Phrenic, vagal and hypoglossal nerve activities (PNA, VNA & HNA, respectively) were first high-pass filtered with a zero-phase 3^rd^ order Butterworth filter (*F*_*c*_ = 300 *Hz*) to remove any DC artifacts before rectification and integration with a moving average filter in forward-backward mode to prevent any phase distortion (*τ* = 100 *ms*). The Kilosort algorithm was used for semi-automated spike sorting of single unit activities recorded on the MEA (37). After spike sorting, we manually inspected and adjusted the cluster assignments. The most frequent modification made to cluster assignments was to remove low-amplitude clusters that were associated with noise or multi-unit activity.

After spike sorting, we sought to assess the distribution of pre-BötC neuronal firing patterns by clustering their respiratory cycle-triggered histograms. We first determined the event times of the inspiratory-to-post-inspiratory (I-PI) phase transition for all breaths via PNA. Depending on the signal-noise ratio of the PNA time series, we used either fixed threshold or the difference between a fast-(*τ* = 33.3 *ms*) and slow- (*τ* = 100 *ms*) moving averages to detect the I-PI transition events. After measuring the average respiratory period, we computed the cycle-triggered average of the respiratory motor pattern and the cycle-triggered histogram of each neurons spiking pattern over 1 respiratory cycle using the I-PI transition events as the trigger for averaging.

To cluster these cycle-triggered histograms of pre-BötC neuronal firing patterns for the group, we combined dimensionality reduction with principal component analysis (PCA) and k-means clustering. The cycle-triggered histograms were scaled to the [0, 1] interval to remove the influence of the peak firing frequencies. Then, we reduced the dimensionality of the scaled group dataset using PCA keeping the top principal components which accounted for >90% of the variance of the original dataset. The inverse transform of these top principal components further illustrated that no meaningful information was lost by discarding the remaining principal components (Suppl. Fig. 2). Then, we determined the optimal number of clusters using the ‘elbow method.’ To visualize the efficacy of the clustering, we examined scaled firing patterns sorted by the k-means cluster labels and used a t-Stochastic Neighbour Embedding to project the dimensionally reduced dataset (and k-means labels) into a 2-dimensional sub-space. Finally, to visualize the firing patterns of each cluster in the respiratory phase domain, we applied the inverse transform of the PCA to each k-means cluster center.

### Population coding & cross-correlation analyses

We first computed the firing rate histogram of each unit in a pre-BötC ensemble recording using a fixed bin width of 50 *ms*. The population rate time histogram was determined for each ensemble by taking the sum of the spike counts of all neurons in an ensemble for all bins (bin width: 50 *ms*) before converting the population counts into the frequency domain. Both the individual pre-BötC firing rate histograms and the population rate were then smoothed with a 2^nd^ order Savitsky-Golay filter with a window length of 5 bins. To assess how the population firing rate or individual neuron firing rates correlated with the three-phase respiratory motor pattern, we measured the Pearson cross-correlation coefficient between VNA and either the population firing rate or the firing rate of an individual pre-BötC neuron. We chose VNA as an index of the three-phase respiratory motor pattern because its pattern reflects all three phases of the respiratory cycle. To test whether the population rate encoded more information about the respiratory motor pattern than individual pre-BötC units, we compared these cross-correlation coefficients using a one-way ANOVA. To further characterize the relative timing between population firing rate peaks and the respiratory motor pattern, we measured the relative time difference between the I-PI transition and the nearest population firing rate peak. Finally, we examined the cycle-triggered averaged respiratory LFP in relation to the respiratory motor pattern and population firing rate.

### Pharmacologic experiments

In all pharmacologic experiments, we spike sorted 10 min of pre-BötC ensemble activity before and after drug administration. In experiments with 8-OH DPAT, neuronal activity was aligned according to the spike templates identified by the Kilosort algorithm. The significance of the increase in respiratory rate was determined using a one-way ANOVA. To assess the effect of 8-OH DPAT on the distribution of pre-BötC firing patterns, the cycle-triggered histograms of all neurons both at baseline and after drug administration were clustered as described above. The distributions of pre-BötC firing patterns were then compared using a two-way Kolmogorov-Smirnov test. In fentanyl experiments, after spike sorting, we used the logISI method to identify bursts and burst-related spikes (38). Once identified, we compared the inter-burst intervals and spikes/burst of baseline bursting, fentanyl-evoked fast- and slow-bursting populations using a one-way ANOVA.

### A Hopfield network model of respiratory pattern generation

We modelled the respiratory pattern generator as a Hopfield network that included fast- and slow-synapses. An all-to-all connected network of *N* = 70 discrete Hopfield neurons was trained via Hebbian learning rules for fast- and slow-synapses to generate a sequential, cyclical pattern of spiking in which various populations were active or silent (29,39,40).

### Network dynamics

Following (29,40), the output of the *i*th neuron, *V*_*i*_(*t*) is related to its net synaptic input *u*_*i*_(*t*) by a gain function *g*[*x*]:

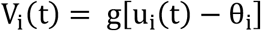

We modelled the neuronal dynamics in the high-gain limit where *g*[*x*] is just the Heaviside step function such that:

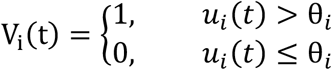

The Hopfield neurons in the model included both fast- and slow-synaptic inputs, 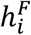 and 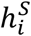 respectively. The net synaptic input to the *i*th neuron, *u*_*i*_(*t*), is:

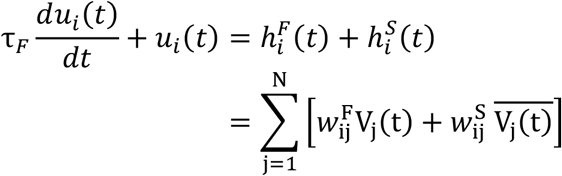

where τ_*F*_ is the time-constant of the fast synapses, 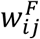 and 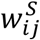 are the synaptic weights of the fast and slow synapses, respectively, and 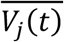 is the time-averaged output of the neuron, i.e.,

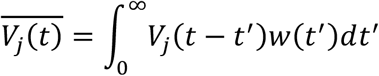

The synaptic response function *w*(*t*) for the slow weights, 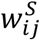, is a non-negative function normalized to unity and characterized by a mean time constant τ_*S*_ satisfying

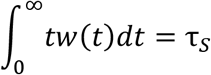

In our model, τ_*F*_ and τ_*S*_ were 5 and 500 timesteps, respectively.

### Hebbian learning of respiratory spiking patterns

The network was trained via Hebbian learning rules to oscillate through a set of states, 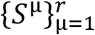, that are each defined by the activity (high-frequency spiking or silent/low-frequency spiking) of all *N* neurons, and that cyclically progress through their defined sequence

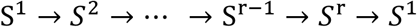

The pre-BötC respiratory firing patterns were not orthogonal (see Fig. 2B for states sequence), and therefore, we followed the method proposed by (40) to encode these non-orthogonal states in the network. We first define the correlation matrix of states as

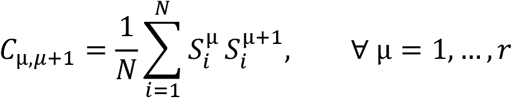

**Figure 2:**
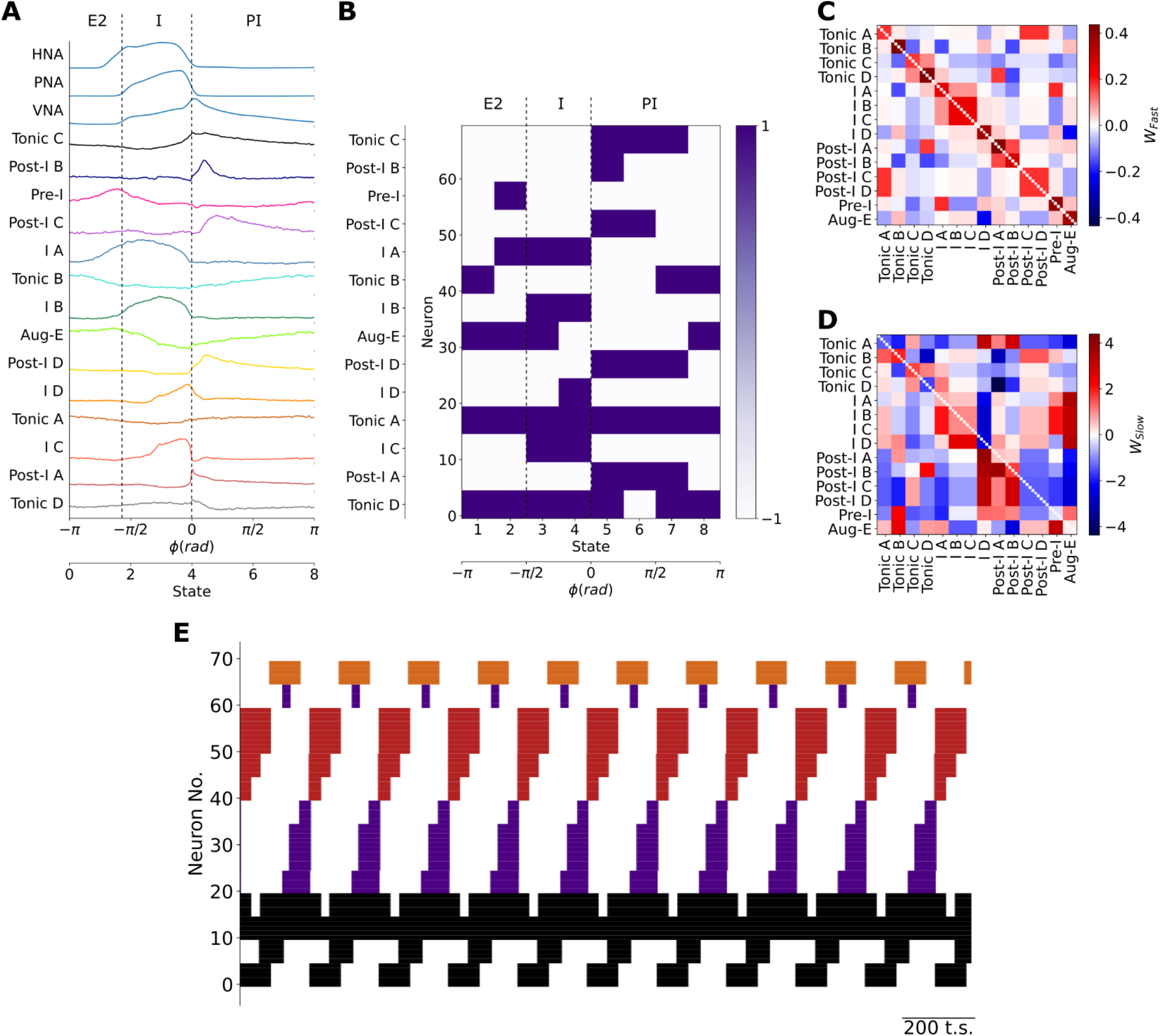
Training a Hopfield model to encode the firing patterns of pre-BötC neuronal clusters. **A** The centroids of pre-BötC neuronal clusters were used as a basis to determine the sequential firing patterns to be encoded in the model. The respiratory cycle was discretized into 8 sequential states to account for the brief firing patterns of the Pre-I, Post-I B and I-D populations. **B** Each pre-BötC cluster was represented by 5 neurons in the model. Their training vectors were taken as +1 when the cluster fired at a high-frequency or -1 when the cluster was silent or firing at a low-frequency. **C, D** The resultant fast-**(C)** and slow-**(D)** synaptic connectivity of the Hopfield network after training to encode the sequential state vectors using Hebbian learning rules. **E** As expected, the model generated the learned sequential firing patterns that underlie the three-phase respiratory motor pattern in the pre-BötC. Black: tonic or respiratory-modulated; Purple: inspiratory; Red: post-inspiratory; Orange: late-expiratory.

Then, orthogonal states can be constructed from linear combinations of the *S*^μ^s

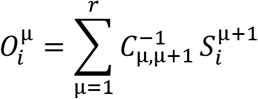

where 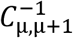 is the pseudo-inverse of the correlation matrix of states.

Finally, the network is trained using the following equations to determine the weights of the fast- and slow-synapses, respectively.

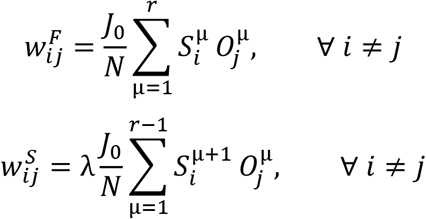

where *J*_0_/*N* sets the scale of the average synaptic strength and λ is a parameter that determines the transition strength between successive states. For all models shown, *J*_0_ was 3, N was 70 and λ was 10.

### Simulations of increasing slow neurotransmission

To simulate the effects of application of a neuromodulatory receptor agonist, we modelled the effects of increasing either slow-inhibition or -excitation in the model by increasing the weights of these synapses by 1.5x their original magnitude.

## Results

### Training a Hopfield network to generate the breathing pattern

Recurrent Hopfield networks that generate periodic activities can be trained via Hebbian learning rules given the rhythmic firing patterns of the network (29,40). Therefore, we first needed to estimate the set of respiratory neuron firing patterns in the intact network of the *in situ* perfused preparation and chose to do so in the pre-BötC since it contains neurons from all three phases of the breathing pattern *in vivo* (30–32).

To accomplish this, we used a silicon high-density MEA to monitor ensemble single-unit activities of the pre-BötC in concert with the respiratory motor pattern on phrenic (PNA), vagal (VNA) and hypoglossal (HNA) motor nerves in the *in situ* perfused brainstem preparation (Fig. 1A & B). We clustered the cycle-triggered histograms of their activity using the transition from inspiration to post-inspiration (I-PI transition) as the trigger for averaging across one respiratory cycle (Fig. 1C-E). Cycle-triggered histograms were determined for 113 neurons from 11 *in situ* preparations (Fig. 1D). To optimize the sensitivity of the clustering to the firing patterns of pre-BötC neurons, the dataset was further pre-processed by scaling to the [0, 1] interval to eliminate the influence of firing rate variability, and by using a principal component analysis (PCA) for dimensionality reduction. After pre-processing, the dataset was clustered using the K-means algorithm (Fig. 1E). The optimal number of pre-BötC neuronal types (k* = 14) was determined using the ‘elbow method’ after repeating the K-means clustering for many values of k.

The pre-BötC of rats *in situ* displayed a mixture of inspiratory, post-inspiratory, late-expiratory and phase-spanning firing patterns (Fig. 1E). As the purpose of our clustering analysis was to develop a consistent, un-biased assessment of the diversity of pre-BötC neuronal types, we avoid introducing a new nomenclature, and instead label the clusters according to a phenotypic division of the classical respiratory neuron types that are often used in central pattern generator models of respiratory pattern generation: pre-inspiratory, inspiratory, post-inspiratory, augmenting-expiratory or tonic/respiratory-modulated. The clustering analysis revealed that of these principal pre-BötC neuronal classes, inspiratory, post-inspiratory and tonic pre-BötC neurons had distinct sub-classes (Fig. 1E). For instance, the clustering analysis revealed that post-inspiratory pre-BötC neurons were further sub-divided into 4 sub-classes (see clusters Post-I A-D, Fig. 1E) that differed in their burst durations and the relative timing of their peak intra-burst frequencies. (Fig. 1F).

To construct the sequential states needed to train the model (see Materials and Methods: Hebbian learning of respiratory spiking patterns), we discretized the firing patterns of each pre-BötC neuronal type (Fig. 2). The cluster centroids of each pre-BötC cluster were taken as the putative firing patterns of each neuronal type (Fig. 2A). The model consists of a network of 70 Hopfield units with fast- and slow-synapses, with each pre-BötC neuronal class represented by 5 model units. The respiratory cycle was sub-divided into 8 sequential states of π/4 radians to enable the approximation of the activity of very transiently active neuron populations like the Post-I B and I D clusters. For each cluster and for each fraction of the respiratory cycle, the state was taken as 1 when the cluster fired at a high frequency and -1 when the population was less active or silent (Fig. 2B). Using these sequential state vectors, we trained the network to encode these sequential firing patterns using Hebbian learning rules. The resultant network weights are shown in Fig. 2C & D. The fast-synapses had a symmetric structure consistent with their role in encoding the fixed points associated with each network state (Fig. 2C), whereas the slow-synapses had an asymmetric structure consistent with their role in destabilizing any given fixed point in the direction of the next sequential fixed point (Fig. 2D). As expected, the trained model generated these sequential firing patterns associated with the three-phase respiratory motor pattern in the absence of external input (Fig. 2E), thereby fulfilling the definition of a central pattern generator network. Thus, we generated an associative memory network model of breathing pattern generation that was constrained by the representation of the three-phase respiratory motor pattern within the pre-BötC. We next validated the predictions of the model experimentally.

### The role of neuromodulation in respiratory rhythm generation

The model encapsulates our hypothesis that the asymmetric connectivity of slow-, neuromodulatory-synapses contributes to respiratory rhythm generation. Because the model contains slow-synapses that are described by their net excitatory or inhibitory effect on the post-synaptic target, we investigated the model predictions associated with this assumption by uniformly increasing the weights of either slow-inhibitory or - excitatory synapses *in silico* (Fig. 3). We chose to increase these slow synaptic weights to enable comparison with the experimental perturbation of systemic administration of neuromodulatory receptor agonists. Increasing the weights of slow-inhibitory synapses in the model evoked only minor effects on network activity (Fig. 3B). Specifically, the sequential firing patterns of the network remained unchanged. However, the frequency of the network’s rhythm increased by 12%. The alternative perturbation, increasing the weights of slow-excitatory synapses, evoked a collapse of the respiratory rhythm (Fig. 3C). With the increase in slow excitatory neuromodulation in the model, the majority of neurons (∼64%) fell silent. The remaining active neurons expressed either tonic or bursting activities. The remaining bursting pattern of activity was characterized by shorter burst durations and inter-burst intervals than any bursting activity observed at baseline (compare Figs. 3A & C). To test these model predictions experimentally, we analyzed the effect of systemic administration of either the 5HT1aR agonist 8-OH DPAT (Fig. 4) or the µ-opioid receptor agonist fentanyl (Fig. 5) on pre-BötC ensemble activity *in situ*.

**Figure 3:**
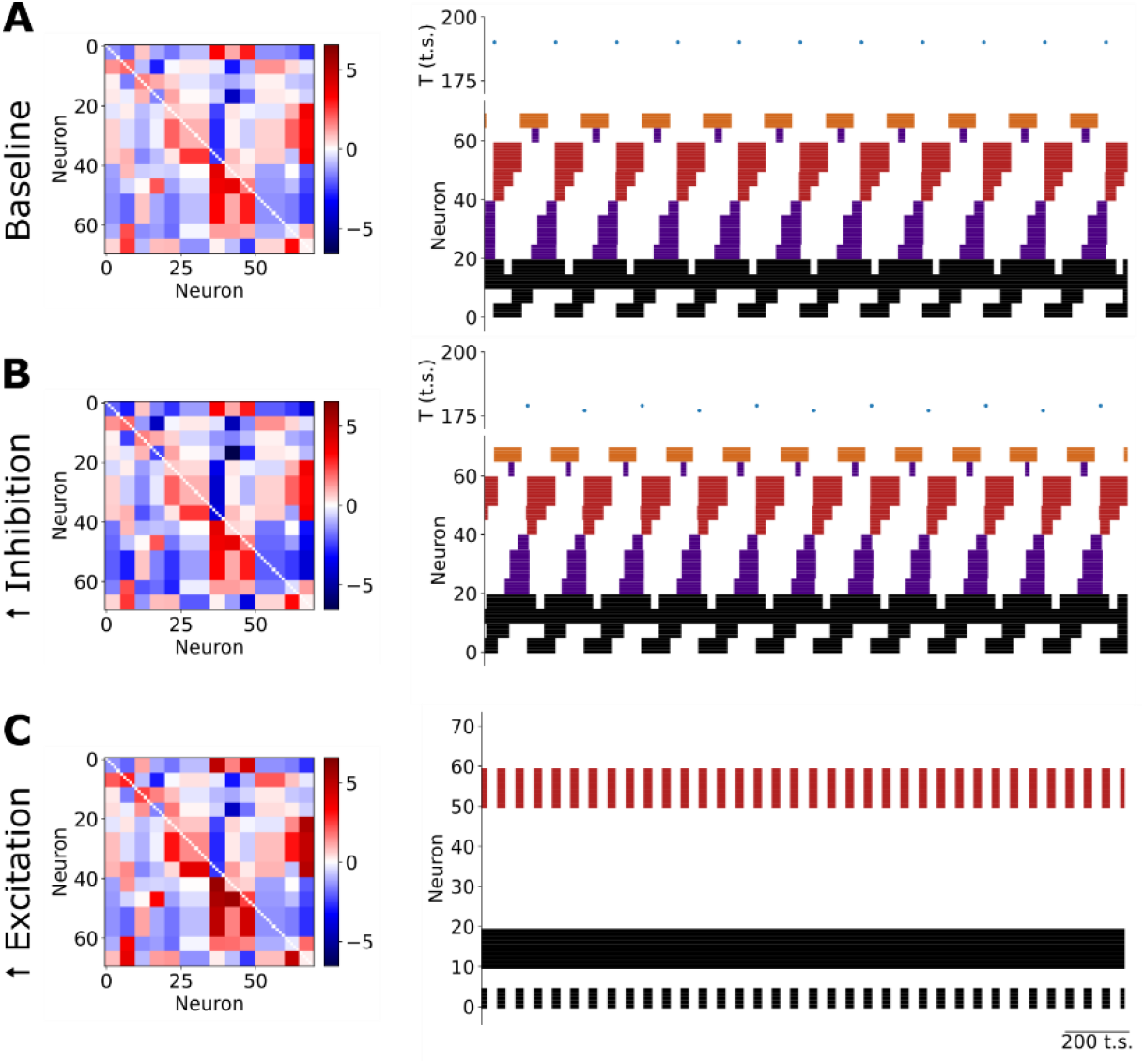
Neuromodulation of the respiratory rhythm *in silico*. **A** The sequential firing pattern produced by the network at baseline. **B** Increasing slow synaptic inhibition in the model evokes an increase in the frequency of the respiratory rhythm without any change in the sequential firing pattern of the network. **C** Increasing slow excitation in the model evokes a collapse of the respiratory rhythm characterized by a reduction in the number of active units whose remaining activity was either tonic or fast bursting.

**Figure 4:**
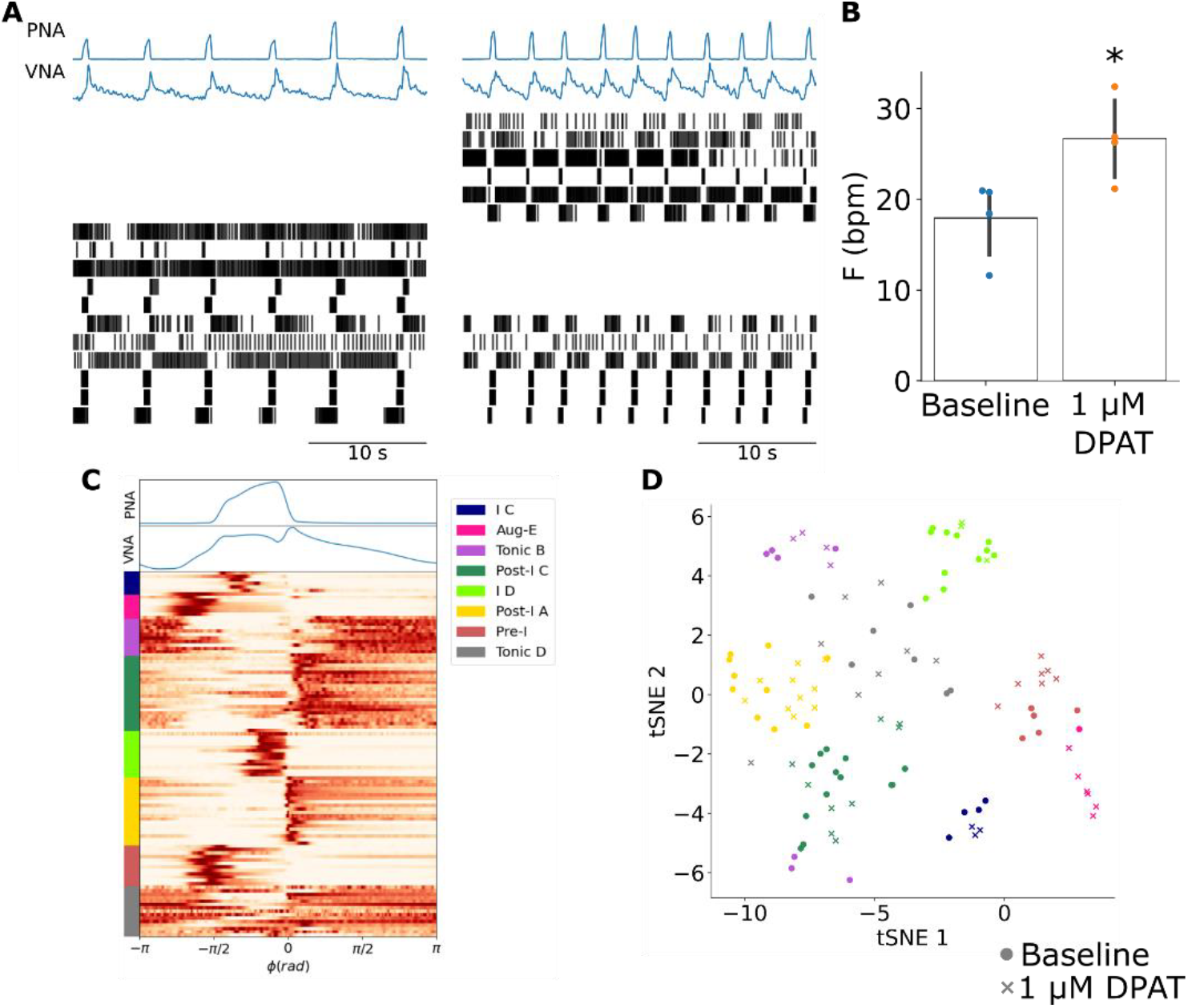
The 5HT1aR agonist 8-OH DPAT increases the frequency of the respiratory rhythm without changing the firing pattern of pre-BötC ensembles. **A** Systemic administration of 8-OH DPAT evoked an increase in the frequency of the respiratory rhythm as observed in PNA & VNA that was associated with a reconfiguration of pre-BötC ensemble activity. In this representative experiment 6 pre-BötC neurons maintained their original firing patterns, but at a higher frequency. In addition, 5 pre-BötC neurons became silent, and 6 pre-BötC neurons were activated. **B** The frequency of the respiratory rhythm was significantly increased after systemic application of 8-OH DPAT. * *p<0*.*05* **C** K-means clustering of all recorded pre-BötC neurons at baseline and after systemic 8-OH DPAT identified many of the pre-BötC neuronal types previously observed. **D** All clusters except the Aug-E population were present at similar ratios at baseline (circles) and after systemic 8-OH DPAT (crosses) suggesting that despite the reconfiguration of pre-BötC ensemble activity, the distribution of neuronal firing patterns that composed the respiratory pattern generator remained the same.

**Figure 5:**
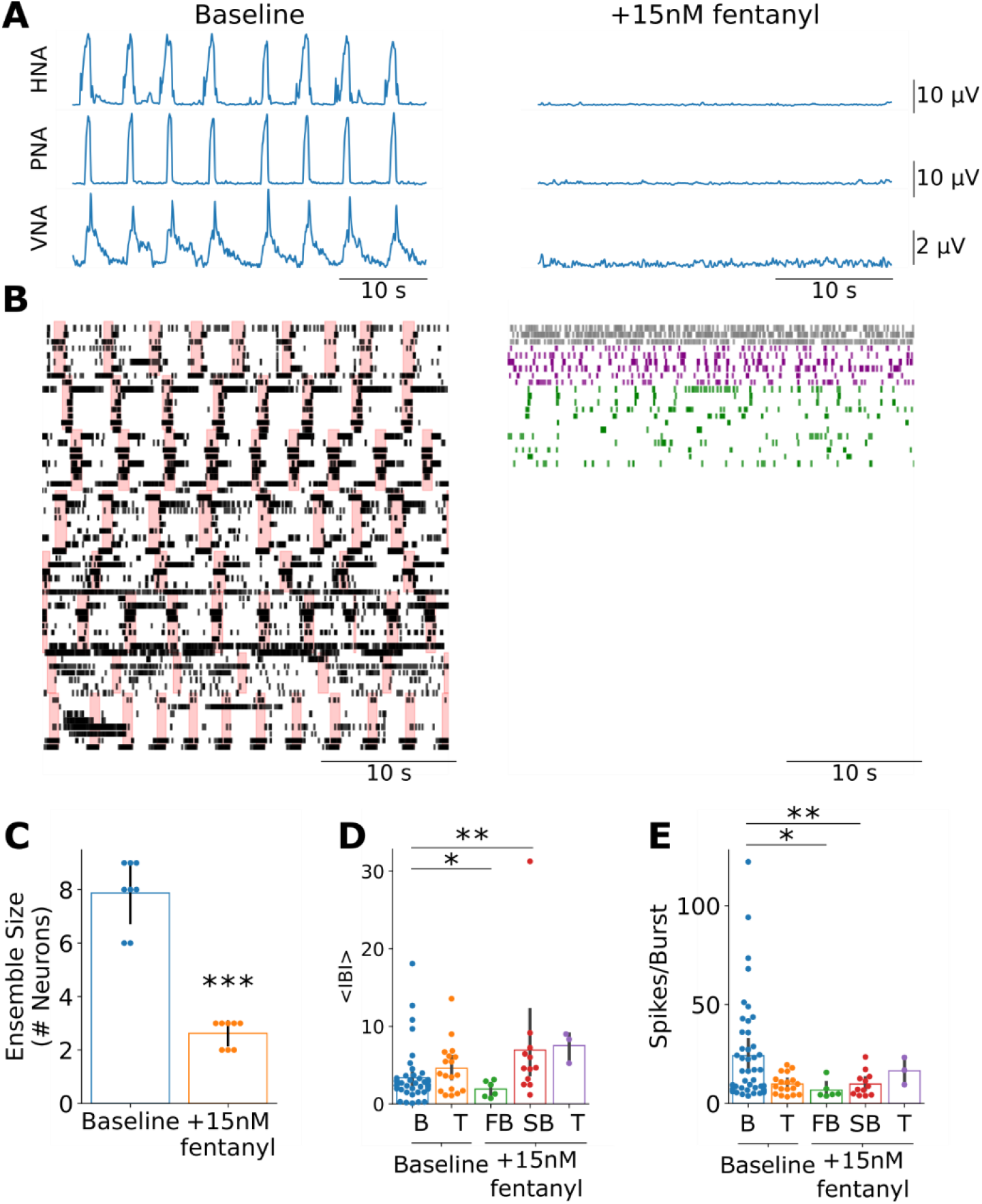
The reconfiguration of pre-BötC ensemble activity after opioid-induced respiratory depression is consistent with model predictions after increasing slow-excitation. **A** Systemic fentanyl administration evokes a collapse of the respiratory rhythm on phrenic (PNA), hypoglossal (HNA) and vagal (VNA) nerves. **B** Consistent with the model predictions after increasing slow excitation, systemic fentanyl administration was associated with a reduction in the size of pre-BötC ensembles and spared tonic (gray), fast-(purple) and slow-(green) bursting firing patterns. Pink bars at baseline indicate the inspiratory periods. **C** Systemic fentanyl administration significantly reduced the number of active neurons in pre-BötC ensembles. *** *p<0*.*001* **D** Consistent with the model predictions, fast-bursting pre-BötC neurons had significantly shorter inter-burst intervals (IBI) than bursting pre-BötC neurons at baseline. However, we also observed a slow-bursting pre-BötC population after systemic fentanyl administration that had significantly longer IBIs than bursting pre-BötC neurons at baseline. B: bursting; T: tonic; FB: fast-bursting; SB: slow-bursting * *p<0*.*05*; ** *p<0*.*01* **E** Consistent with the model predictions both fast- and slow-bursting pre-BötC neurons fired fewer spikes per burst than bursting pre-BötC neurons at baseline. * *p<0*.*05*; ** *p<0*.*01*

Systemic application of 8-OH DPAT evoked effects on pre-BötC ensemble activity that were qualitatively similar to the model predictions of increasing slow inhibition. 8-OH DPAT increased the frequency of the respiratory rhythm (Fig. 4A & B). This increase in respiratory rate was accompanied by a reconfiguration of pre-BötC ensemble activity wherein some units became silent, previously silent units became active and some units maintained their baseline firing patterns (Fig. 4A). To test whether the distribution of respiratory firing patterns was altered, we clustered the cycle-triggered histograms of all units before and after systemic 8-OH DPAT (Fig. 4C). All firing pattern clusters contained units from both baseline and 8-OH DPAT groups (Fig. 4D). Importantly, the distribution of pre-BötC firing patterns after systemic 8-OH DPAT was not significantly different from that at baseline (*p=0*.*98*). Taken together, these results suggest that exogenous enhancement of 5HT1aR transmission evokes qualitatively similar effects as those predicted by an increase of slow inhibition in the model.

The experimentally observed effects of fentanyl on pre-BötC ensemble activity were qualitatively similar to the model predictions of increasing slow excitation. Systemic administration of 15 nM fentanyl evoked a collapse of the respiratory motor pattern on phrenic, vagal and hypoglossal nerves (Fig. 5A).

Consistent with the model, ensemble activity in the pre-BötC was largely suppressed with the number of active neurons from 7.8 ± 1.2 to 2.6 ± 0.5 neurons (Fig. 5B & C, *p<0*.*001*). Further, pre-BötC neuronal activity after systemic fentanyl administration consisted of neurons with either tonic or bursting activities. However, unlike the model, we observed bursting neurons with either fast-bursting and slow-bursting phenotypes. Consistent with the model, both classes of bursting neurons had shorter burst durations than at baseline, firing significantly fewer spikes per burst (Fig. 5E, Fast-Bursting: *p<0*.*05*, Slow-Bursting: *p<0*.*01*). Further, like the model predictions, the fast-bursting class also had significantly shorter inter-burst intervals than bursting pre-BötC neurons at baseline (Fig. 5D, *p<0*.*05*). However, the slow-bursting class had significantly longer inter-burst intervals than bursting pre-BötC neurons at baseline (Fig. 5D, *p<0*.*01*). Taken together, the model predictions after perturbing neuromodulatory transmission were consistent with experimental results suggesting that neuromodulation underlies respiratory rhythm generation.

### Population activity encodes respiratory phase transitions

Another prediction of the model was the existence of a population code of the respiratory motor pattern (Fig. 6). In the model, transitions between successive states occur because of the slow-synaptic neurotransmission that changes the energy landscape of the network. During any given state, the network lies at a global energy minimum leading to the repetitive firing of neurons associated with that fixed point. Because of the asymmetric connectivity of the slow synapses in the network, each fixed point is progressively destabilized until the fixed point associated with the next sequential state becomes the new global minimum. When this critical point is reached, the network rapidly transitions to the new fixed point thereby recalling the activity pattern of the next sequential state. These transitions are not instantaneous. Each state transition involves a short overlap of the activity patterns associated with the two successive states during this pattern recall process causing peaks in the population firing rate. We observed that the population activity of the network resembled the bi-phasic waveform expressed in vagal nerve activity, and that three of the eight state transitions—from I to PI, from PI to E2 and from E2 to I—were associated with brief peaks in population activity (Fig. 6A, *top*) when the distance between successive state vectors was maximal.

**Figure 6:**
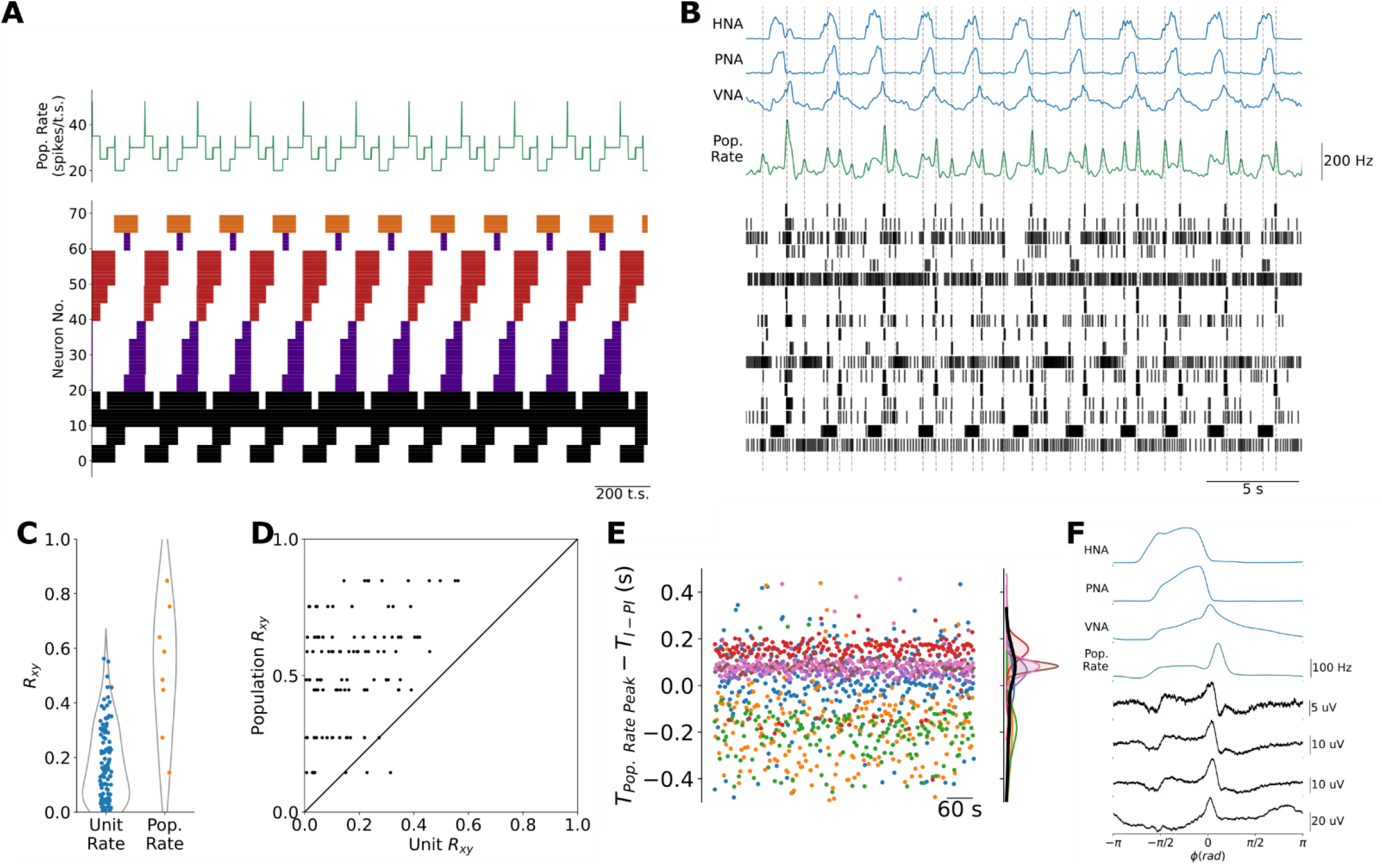
Population coding of the transitions between respiratory phases in the model and in the pre-BötC *in situ*. **A** Because the recall of the next sequential state involves a slight overlap of sequential assemblies, the model generates brief peaks in the population firing rate at each of the three transitions between respiratory phases when the adjacent state vectors are most distant. Black: tonic or respiratory-modulated; Purple: inspiratory; Red: post-inspiratory; Orange: late-expiratory. **B** Consistent with the model predictions, pre-BötC ensemble activity is associated with a population firing rate that also peaks at or near the transitions between respiratory phases. **C** The cross-correlation between the population firing rate and the three-phase respiratory pattern of vagal nerve activity (VNA) was significantly greater than that between any individual pre-BötC unit and VNA. *** *p<0*.*001* **D** The cross-correlation between the population firing rate and VNA was almost always greater than that between the unit firing rate and VNA. KS-test: p<0.001. **E** Individual pre-BötC ensembles varied with respect to the precision with which their population firing rate encoded the I-PI transition. **F** Cycle-triggered averages of pre-BötC local field potentials (LFPs) more reliably encoded the transitions between respiratory phases.

To test whether the intact respiratory network *in situ* also generates a population code of respiratory phase transitions, we measured the population firing rate of pre-BötC ensembles *in situ*. Like the model, the population activity of the pre-BötC ensembles was characterized by a basal level of activity interspersed with brief peaks of fast spiking (Fig. 6B). Consistent with the model, the peaks in population activity occurred at or near the three transitions between respiratory phases. The cross-correlation between the population activity and the vagal motor pattern, which carries information about all three phases of the respiratory motor pattern, was significantly greater than that of individual neurons (Fig. 6C) suggesting that the population activity carries more information about the breath-by-breath respiratory motor pattern than the activity of any individual pre-BötC neuronal type.

To further investigate whether the timing of pre-BötC population activity was specifically associated with the transitions between respiratory phases, we focused on the relative timing between ensemble population activity peaks that occurred nearest to the transition between I and PI (Fig. 6D). On average, the ensemble population activity peak occurred 0.0024 ± 0.206 s before the decline in inspiratory PNA amplitude reflecting the fact that individual ensembles differed greatly with respect to the precision of and relative timing of their encoding the I-PI transition (Fig. 6D). We hypothesized that this variability may be due to the limited number of pre-BötC neurons that we were able to simultaneously monitor using silicon MEAs. Therefore, we further addressed this question by measuring the cycle-triggered averages of respiratory local field potentials (LFPs) on each of the four MEA shanks and ensemble population activity in a representative experiment since LFPs reflect the synaptic activity of many more neurons (Fig. 6E). Respiratory LFPs in the pre-BötC occurred specifically at the E2-I and I-PI transitions, whereas the ventral-most site of the fourth shank identified respiratory LFPs occurring specifically at the I-PI and PI-E2 transitions. Taken together, these data confirm the prediction of the model that population activity within the respiratory network peaks at the transitions between respiratory phases, a feature that is not present in the population activity of CPG models of respiratory pattern generation (Suppl. Fig. 3).

## Discussion

In this study, we have developed a Hopfield network model of respiratory rhythm and pattern generation that encapsulates the hypothesis that slow-, neuromodulatory-connectivity in the respiratory network is organized asymmetrically to generate the respiratory rhythm. We tested this model assumption by comparing the predictions of uniformly increasing slow-inhibitory or -excitatory weights with *in situ* experiments in which we recorded ensemble activity of the pre-BötC before and after systemic administration of 5HT1aR or µ-OR agonists. Increasing slow-inhibitory weights in the model or activating 5HT1aRs systemically with 8-OH DPAT increased the frequency of the respiratory rhythm without changing the firing patterns of respiratory neurons. Increasing slow-excitatory weights in the model or activating µ-ORs systemically with fentanyl arrested the respiratory rhythm sparing neurons with tonic- and short bursting-firing patterns. The similarity between model predictions and experiments supports the hypothesis that neuromodulatory connectivity in the respiratory network is organized asymmetrically to promote rhythmogenesis. The model also predicted the existence of a population code of respiratory phase transitions which we confirmed in the population activity of pre-BötC ensembles and respiratory LFPs in the pre-BötC.

### Network models of respiratory rhythm and pattern generation

Computational models of respiratory rhythm and pattern generation have been developed to explain experimental observations at both cellular and network scales. The discovery of spontaneously bursting pre-inspiratory neurons of the pre-BötC led to the development of cellular models that describe how persistent sodium currents could underlie spontaneous inspiratory bursting in single neurons (41). Recent work on the bursting mechanisms of the isolated pre-BötC has highlighted that its small-world connectivity, rather than its intrinsic conductances, underlies the capacity to generate inspiratory bursting activity (42,43). This conceptual model has been incorporated into network models that consist of excitatory neurons containing a subset of spontaneously bursting units connected in a small-world pattern, which is now considered to explain the inspiratory bursting of the isolated pre-BötC (44,45). Beyond the pre-BötC inspiratory activity, several network models have been developed to formalize the long-standing conceptual view of the respiratory network as a central pattern generator (46–51). These central pattern generator models consist of neurons with mutual inhibitory interactions that sculpt respiratory neuronal activities from several sources of excitatory drive. Importantly, they have shown that reciprocal inhibition can account for a variety of experimental observations including the generation of the three-phase respiratory motor pattern of inspiration, post-inspiration and late-expiration. However, a limitation of previous models is that the respiratory network is not composed of strictly inhibitory or excitatory neurons. For the case of central pattern generator models, this property is highlighted by studies which show that blockade of synaptic inhibition in key inspiratory or expiratory compartments of the respiratory network is not sufficient to ablate the respiratory pattern *in vivo* (9,52). Thus, there remains a need for computational models of respiratory rhythm and pattern formation that have greater face validity.

In the present study, we developed a network model of respiratory rhythmogenesis that incorporated excitatory, inhibitory and neuromodulatory connections. Using previously proposed connectivity patterns (29,40), we were able to generate a network model of respiratory rhythm and pattern generation based on the assumed set of respiratory firing patterns which we measured from ensemble recordings of the pre-BötC *in situ*. This model encapsulated our hypothesis that asymmetric neuromodulatory connections can promote the generation of the respiratory rhythm. In testing this core assumption, we found that the model was also predictive in that perturbations of its connectivity weights were consistent with experimental perturbations of serotonergic or opioidergic neurotransmission. However, the model is not without limitation. For instance, the model does not include spontaneous bursting neurons. However, spontaneous bursting neurons have been shown to be dispensable in central pattern generator models of the respiratory rhythm (53). Further, the Hopfield units of our model are binary and thus cannot generate the spiking or bursting dynamics associated with more detailed neuron models. However, it was demonstrated that the network connectivity patterns of Hopfield networks can be translated into those for spiking networks to yield networks with similar behavior (54). Finally, the present model also does not follow Dale’s law since the Hopfield units can have both excitatory and inhibitory neurons. Given such significant simplifications from the biological system, it is notable that the model was able to predict the collapse of network activity following fentanyl administration.

### Population coding of respiratory phase transitions

The findings of the present study also extend previous observations of a population code of respiratory phase transitions. In an earlier study, we reported that respiratory LFPs, which reflect the synaptic activities of local populations, peaked specifically at the transitions between the three phases of the respiratory cycle throughout the ponto-medullary respiratory network (55). In the present study, this feature was observed both in the model in the ensemble activity of the pre-BotC *in situ*. In the model, transitions between states are evoked by slow neuromodulatory transmission that acts to change the ‘energy’ landscape of the network such that the fixed point associated with a given state is destabilized in the direction of the fixed point associated with the next state (40). At the transition, a partial cue of the next state’s memory is established allowing the network to recall the activity pattern of the next state. These transitions involve a brief overlap of the activities of adjacent Hebbian assemblies as the memory of the next fixed point is recalled and stabilized. Interestingly, peaks in the network’s population activity appeared at only three of eight transitions, which corresponded to those between inspiration, post-inspiration and late-expiration. We observed a qualitatively similar pattern of population activity in pre-BötC ensembles *in situ*. Importantly, this property of population activity cannot be accounted for in half-centre oscillator CPG models of respiratory pattern generation (Suppl. Fig. 3) since transitions in such models occur via an escape or release mechanism in which the population activity at a transition is either balanced or shifts to a new plateau depending on the number of units active before and after the transition (49). Thus, the proposed model of asymmetric neuromodulation better accounts for the population activity of the respiratory network *in situ*.

In addition, we observed, both in the model and in experiments, that the cycle-triggered average of the population activity in the network or in the pre-BötC respectively resembled the bi-phasic discharge of the vagal motor pattern, which regulates upper-airway patency. This observation is consistent with the recent characterization of a role for the pre-BötC in regulating, not just inspiratory discharge in the inspiratory motor nerves, but also in the inspiratory and post-inspiratory activity in the vagus (56). In the model, this bi-phasic pattern of population activity reflects the distribution of firing patterns the network was trained to generate. We derived this distribution directly from the clustering of firing patterns present in ensemble recordings *in situ*.

Consistent with previous observations in the intact brainstem, the distribution of pre-BötC firing patterns included neurons with bursting activity in the inspiratory, post-inspiratory and late-expiratory phases as well as neurons with tonic or respiratory modulated activities (30–32). The latter classes of respiratory neurons have been previously implicated in respiratory phase switching and the reflex and behavioral control of the respiratory pattern (57–60). In contrast, in our model, these patterns of activity are merely a consequence of the overall network connectivity, with each population’s slow synapses playing significant roles in determining respiratory phase switching. Consistent with this experimental finding, we observed a stronger cross-correlation between pre-BötC ensemble activity and the vagal motor pattern than the activity of any individual pre-BötC neuron suggesting that the population, rather than individual bursting neurons, is responsible for encoding the respiratory motor pattern in network activity. Together, these experimental data confirm the model prediction of the temporal structure of population activity within the respiratory network.

### Implications for opioid-induced respiratory depression (OIRD)

OIRD remains a significant health problem in the United States (12,61). Recently, the risk posed by illicit synthetic µ-OR agonists has been further exacerbated by the presence of adulterants like xylazine that act on α2 adrenergic receptors and nitazenes which are µ-OR agonists that may not be fully counteracted by the µ-OR antagonist naloxone (61). Thus, the incidence of OIRD due to synthetic opioids and combinations of opioid and non-opioid substances has motivated the need to discover new therapeutics to counteract OIRD. Our computational model and experimental results suggest that neuromodulatory connectivity within the respiratory network is organized asymmetrically to promote rhythmogenesis. We propose that the pattern of neuromodulation should be considered for the rational design of therapies to treat respiratory disorders like OIRD. More specifically, our results suggest that identification of alternative neuromodulatory targets to prevent or reverse OIRD will require the consideration of the pattern of neuromodulator receptor expression, its overlap with that of µ-OR expression and the firing pattern of the target respiratory neurons.

Neuromodulatory signaling pathways have long been therapeutic targets for respiratory disorders. A remarkable example of this strategy occurred in the case of a patient who experienced severe apneustic respiratory disturbances after surgical resection of a tumor at the ponto-medullary junction (62). In this case study, the apneustic respiratory motor pattern was corrected without side-effects using buspirone, a 5HT1aR agonist. The rationale behind the therapy arose from the perspective that neuromodulators act to influence intracellular second messenger cascades and that counteracting the influence of one pathway could be achieved by activating alternative second messenger systems with the right neuromodulatory agonist (63). However, the predicted downstream effects on membrane excitability came from intracellular recordings of very few respiratory neurons before and after drug applications. This limited evidence also led to other cases in which neuromodulatory therapies were met with limited success. For instance, 5HT4aRs were identified as a therapeutic to better manage OIRD without the loss of analgesia that accompanies OIRD reversal with naloxone (64). However, later clinical trials showed that the 5HT4R-agonist mosapride was ineffective in recovering from morphine-induced OIRD in humans (65). In the case of the irregular respiratory rhythms present in patients with Rett syndrome, pre-clinical studies in MeCP2-deficient mice developed strong evidence that drugs targeting serotonergic and dopaminergic receptors were effective to correct respiratory disturbances (66,67). However, clinical trials in Rett patients treated with saritozan, a 5HT1aR- and D2R-agonist, were unsuccessful (68).

In the present study, we developed a computational model of respiratory pattern generation based on the hypothesis that the pattern of neuromodulation in the network is organized asymmetrically to promote periodic sequential network activity. Despite the simplicity of this network model, increasing the weights of slow-excitatory neuromodulatory connections accurately predicted the pattern of network activity that was experimentally observed following fentanyl-induced OIRD, specifically a reduction in the number of active neurons that spared populations with either tonic or short bursting activities. Similarly, increasing the weights of slow-inhibitory neuromodulatory connections predicted the effects of systemic application of 5HT1aR agonist 8-OH DPAT which increases the frequency of the respiratory rhythm without changing the pattern of respiratory network activity. This latter model prediction has been widely observed in pharmacologic, optogenetic or chemogenetic experiments both in reduced slice preparations *in vitro* and in the intact network *in situ* for many neuromodulatory systems (6) including, for example, serotonin (8,64,69), acetylcholine (70), norepinephrine (71,72), dopamine (73), ATP (74–76) and histamine (77). That a relatively simple model of respiratory pattern generation could predict the effects of neuromodulation highlights the need to consider the pattern of neuromodulation across the network for the rational design of neuromodulatory therapies. In other words, one should address the question of whether a proposed neuromodulatory therapeutic targets the opposing asymmetric respiratory neuronal populations to promote respiratory pattern formation? Nonetheless, these simulations and experiments support previous suggestions to develop combinatorial neuromodulatory therapies, particularly to protect against opioid-induced respiratory depression (8,64,78).

The need to consider the network mechanism of respiratory neuromodulation is further highlighted by the fact that both neuromodulatory agonists used in the present study are coupled to G_i/o_-dependent signaling cascades (8,79). In the case of the 5-HT1aRs, our findings were consistent with a predominant effect of slow-inhibition in the network. In the case of µ-ORs, our results which suggest a net effect of opioids to increase slow-excitation may appear counter-intuitive to the commonly held notion that activation of µ-ORs evokes inhibition of membrane excitability. Importantly, the action of a particular neuromodulatory receptor agonist on one cell-type does not necessarily generalize to its effect on any neuron. Instead, the effect of activating neuromodulatory receptors depends on the targets of the corresponding intracellular signaling cascades which vary across neuronal cell types. In the case of µ-ORs, it is well known that neurons can show either excitatory or inhibitory effects depending on the cell-type (79). One simple explanation of our observations is that µ-ORs may have a greater effect on inhibitory neurons such that the net effect of opioids at the level of the respiratory network is that of a slow-disinhibition. Alternatively, it has also been shown that µ-ORs can directly excite their target neurons via the coupling of their G_βγ_-subunits to PLC-dependent signaling cascades that increase intracellular calcium levels (80). In either case, our computational and experimental observations supporting an asymmetric pattern of neuromodulation in the network further highlight the need to consider the intracellular effects of a neuromodulatory pathway across the whole network, rather than in small subsets of respiratory neurons.

## Acknowledgements

This study was supported by NIH R01 HL161582 (MD & RRD), NIH R01 HD111415 (PMM), NIH RF1 NS118606-01 (PJT), NSF DMS-2052109 (PJT), the Oberlin College Department of Mathematics (PJT), the HRC (JFRP), the Royal Society Te Apārangi (JFRP) and the Sidney Taylor Trust (JFRP).

