## Supplementary Information for "Asymmetric neuromodulation in the respiratory network contributes to rhythm and pattern generation"

### Supplementary Methods

#### Registration of neuronal locations in the Waxholm atlas coordinate space

In a subset of experiments, both to confirm the localization of these neuronal types to the pre-BötC and to assess their spatial distribution within the pre-BötC, we developed an approach to determine the rigid transformation that would register the coordinate space of the MEA micro-positioner to the Waxholm 7T MRI atlas of the Sprague-Dawley rat brain (36). To do so, we first measured the coordinates of 5 brainstem surface landmarks of the floor of the 4$^{th}$ ventricle that are readily observable in both the *in situ* preparation and in the Waxholm atlas (Suppl. Fig. 1). Using these correspondence points, we calculated the rigid transformation that would register these two coordinate spaces using the analytic method proposed by (81) (Suppl. Fig. 1b). After registering the MEA recording sites to the Waxholm MRI atlas of the Sprague-Dawley rat brain, we estimated the 'centre-of-mass' of each neuron's spike template to determine the location of each neuron relative to the MEA. Then, we applied the rigid transformation to these MEA unit location coordinates to determine the anatomic location of each neuron within the Waxholm atlas. This method enabled an estimation of the position of the MEA electrode sites within the Waxholm atlas and confirmed that the recorded ensembles were localized to the pre-BötC (Fig. 1a & f).

#### Respiratory CPG model

To illustrate the population activity associated with half-centre oscillator models of respiratory pattern generation, we simulated the respiratory CPG model of (49). The model consists of four neurons representing pre-I, early-I, post-I and aug-E populations. These neuronal populations were connected via reciprocal inhibition and received common excitatory drive from three sources. The membrane potential of the pre-I neuron $V_1$ is intrinsically bursting due to its persistent sodium current and was defined by the following equation:

$$\mathrm{C}\frac{dV_1}{dt} = -\mathrm{I_{NaP}} - \mathrm{I_K} - \mathrm{I_{L_1}} - \mathrm{I_{SynE_1}} - \mathrm{I_{SynI_1}}$$

where $I_{NaP}$ is the persistent sodium current and $I_K$ is the delayed rectifier potassium current.

The membrane potential of the other neurons ($i \in 2,3,4$) was

$$C\frac{dV_i}{dt} = -I_{AD_i} - I_{L_i} - I_{SynE_i} - I_{SynI_i}$$

where $I_{AD_i}$ is an outward potassium current that mediates adaptation behaviour. In both of the above equations, $C$ is the membrane capacitance, $I_{L_i}$ are the leak currents, $I_{SynE_i}$ and $I_{SynI_i}$ are the excitatory and inhibitory synaptic currents.

The membrane currents are defined by the following equations

$$I_{NaP} = \overline{g}_{NaP} m_{NaP} h_{NaP} (V_1 - E_{Na})$$

$$I_K = \overline{g}_K m_K^4 (V_1 - E_K)$$

$$I_{AD_i} = \overline{g}_{AD} m_{AD_i} (V_i - E_K)$$

$$I_{L_i} = \overline{g}_L (V_i - E_L)$$

$$I_{SynE_i} = \overline{g}_{SynE}\left(V_i - E_{SynE}\right)\sum_{k=1}^{3} c_{ki}\, d_k \quad \forall\ i \neq 2$$

$$I_{SynE_2} = \overline{g}_{SynE}\left(V_2 - E_{SynE}\right)\left[a_{12} f_1(V_1) + \sum_{k=1}^{3} c_{ki}\, d_k\right]$$

$$I_{SynI} = \overline{g}_{SynI}\left(V_i - E_{SynI}\right)\sum_{j=2}^{4} b_{ji}\, f_j\left(V_j\right) \quad \forall\ j \neq i$$

where $\overline{g}_{NaP}$, $\overline{g}_{K}$, $\overline{g}_{AD}$, $\overline{g}_{L}$, $\overline{g}_{SynE}$ and $\overline{g}_{SynI}$ are the maximal conductances of the corresponding currents, $E_{Na}$, $E_K$, $E_{AD}$, $E_L$, $E_{SynE}$ and $E_{SynI}$ are the corresponding reversal potentials, $a_{12}$ is the synaptic weight from the pre-I to early-I neuron, $b_{ji}$ is the inhibitory synaptic weight from neuron $j$ to neuron $i$, and $c_{ki}$ is the weight of the synaptic drive from drive $d_k$ ($k \in 1,2,3$) to neuron $i$.

The nonlinear function $f_i(V_i)$ defines the spiking activity of each neuron

$$\mathrm{f_i(V_i)} = \frac{1}{1 + \mathrm{e}^{\frac{-(\mathrm{V_i} - \mathrm{V_{1/2}})}{\mathrm{k_{V_i}}}}} \quad \forall\ \mathrm{i} \in 1,2,3,4$$

where $V_{1/2}$ is the half-activity voltage and $k_{V_i}$ defines the slope of output function of each neuron.

The slow inactivation of the persistent sodium current is

$$\tau_{h_{NaP}}(V_1)\frac{d}{dt}h_{NaP} = h_{\infty NaP}(V_1) - h_{NaP}$$

The slow adaptation of the other three neurons are

$$\tau_{AD_i}\frac{d}{dt}m_{AD_i} = k_{AD_i} f_i(V_i) - m_{AD_i}$$

where $\tau_{AD_i}$ is a fixed time constant and $k_{AD_i}$ is the maximal adaptation.

The voltage-dependent activation and inactivation variables and time constant for the persistent sodium and rectifying potassium channels of neuron 1 are

$$m_{NaP} = \frac{1}{1 + e^{-(V_1+40)/6}}$$

$$\mathrm{h_{\infty NaP}} = \frac{1}{1 + \mathrm{e}^{(\mathrm{V_1}+48)/6}}$$

$$\tau_{h_{NaP}} = \frac{\tau_{h_{NaP}max}}{cosh(V_1 + 48)/12}$$

$$m_K = \frac{1}{1 + e^{-(V_1+29)/4}}$$

Model parameters are defined in Supplementary Table 1. The model was numerically integrated using the fourth-order Runge-Kutta method with a timestep size of 1 ms. After simulation, we computed the population firing rate by summing the firing rates of the neurons, $f_i(V_i)$, and scaling by the network size, $n = 4$.

**Supplementary Table 1: Respiratory CPG model parameters.**

| **Parameter (unit)** | **Values** |
|---|---|
| Membrane capacitance (pF) | $C = 20$ |

| | |
|---|---|
| Maximal conductances (nS) | $\overline{g}_{NaP} = 5.0$, $\overline{g}_{K} = 5.0$, $\overline{g}_{AD} = 10.0$, $\overline{g}_{L} = 2.8$, $\overline{g}_{SynE} = 10.0$ and $\overline{g}_{SynI} = 60.0$ |
| Reversal potentials (mV) | $E_{Na} = 50$, $E_{K} = -85$, $E_{AD} = -60$, $E_{L} = -60$, $E_{SynE} = 0$ and $E_{SynI} = -75$ |
| Synaptic weights | $a_{12} = 0.4$, $b_{21} = 0$, $b_{23} = 0.25$, $b_{24} = 0.35$, $b_{31} = 0.3$, $b_{32} = 0.05$, $b_{34} = 0.35$, $b_{41} = 0.2$, $b_{42} = 0.35$, $b_{43} = 0.1$, $c_{11} = 0.115$, $c_{12} = 0.3$, $c_{13} = 0.63$, $c_{14} = 0.33$, $c_{21} = 0.07$, $c_{22} = 0.3$, $c_{23} = 0$, $c_{24} = 0.4$, $c_{31} = 0.025$, $c_{32} = c_{33} = c_{34} = 0$ |
| Parameters of spiking activity functions (mV) | $V_{1/2} = -30$, $k_{V_1} = 8$, $k_{V_2} = k_{V_3} = k_{V_4} = 4$ |
| Time constants (ms) | $\tau_{h_{NaP}max} = 6000$, $\tau_{AD_2} = \tau_{AD_4} = 2000$, $\tau_{AD_3} = 1000$ |
| Adaptation parameters | $k_{AD_2} = k_{AD_4} = 0.9$, $k_{AD_3} = 1.3$ |

**Supplementary Figures**

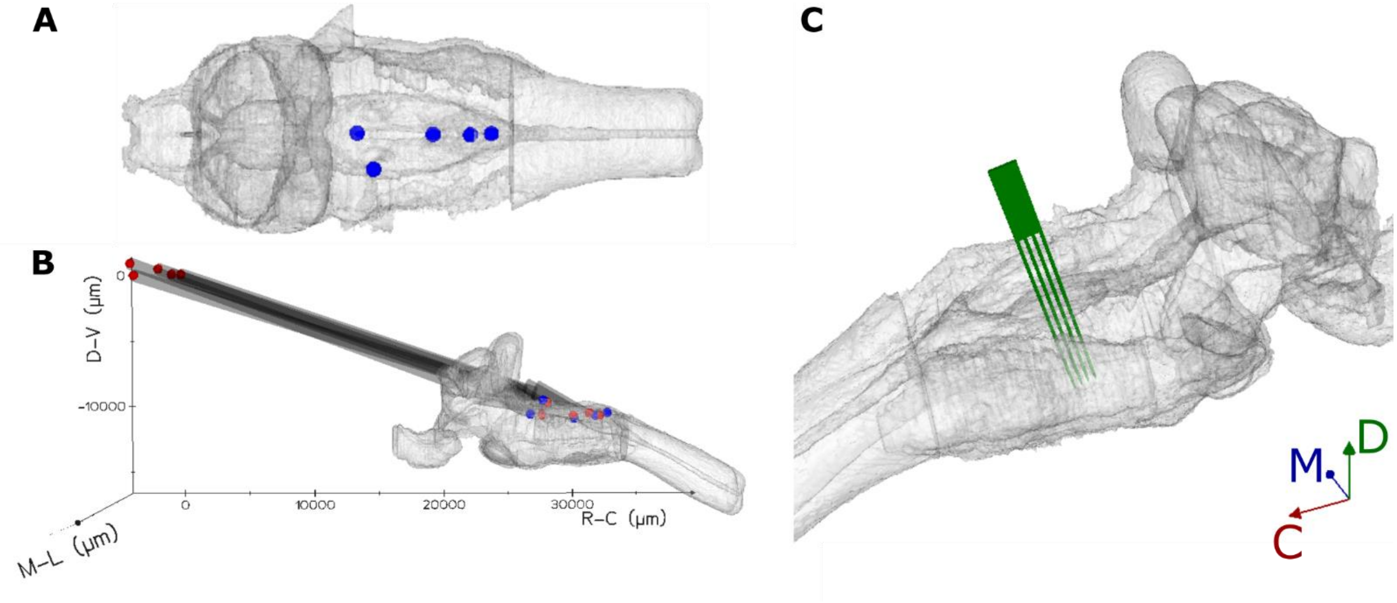


**Supplementary Figure 1: Registration of MEA coordinate space to Waxholm atlas of Sprague-Dawley rat brain.**

**A** Locations of brainstem surface landmarks in the Waxholm atlas of the Sprague-Dawley rat brain.

**B** Registration of the observed surface landmark coordinates (red) to those of the Waxholm atlas (blue).

**C** Reconstructed MEA positioning in a representative experiment.

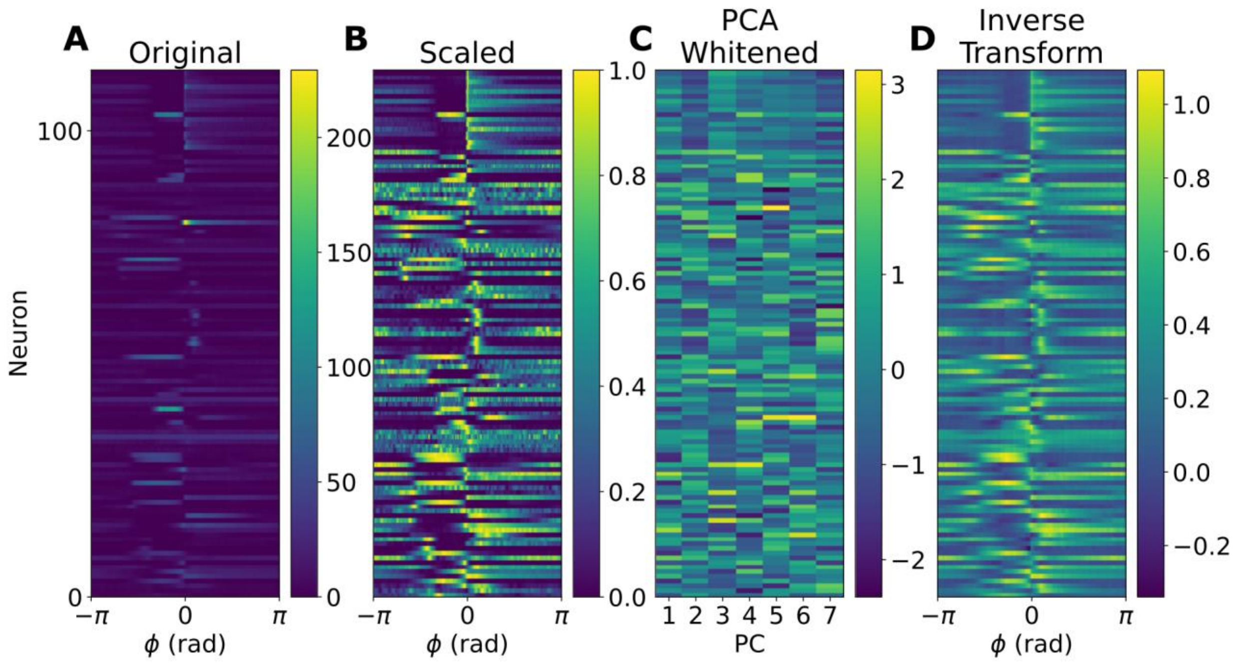


**Supplementary Figure 2: Pre-processing of cycle-triggered histograms for k-means clustering.**

**A** Cycle-triggered histograms (CTHs) for all pre-BötC neurons in Hz.

**B** Scaled CTHs for all pre-BötC neurons in a.u.

**C** The dimensionality of the dataset was reduced with a PCA keeping the top 7 components which accounted for ~90% of the original variance.

**D** Inverse transform of the dimensionality reduced dataset shows that no meaningful information about the cycle-triggered firing rate patterns was lost by discarding the bottom principal components.

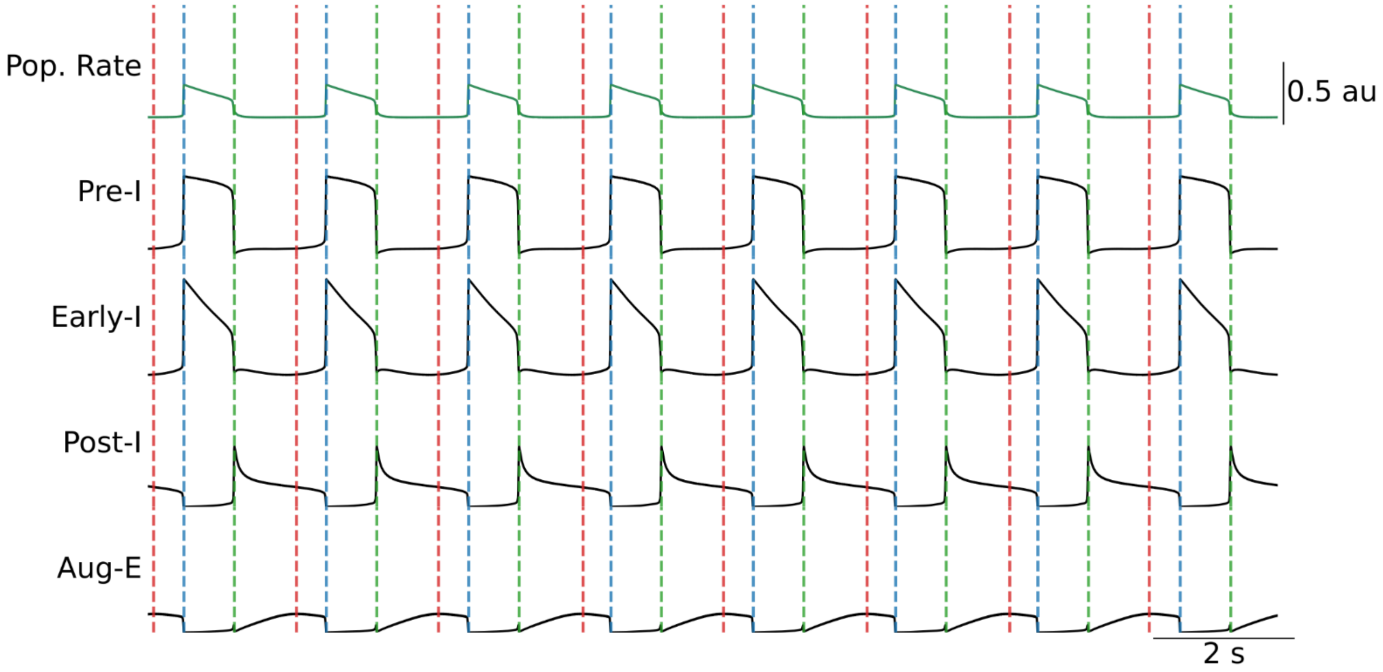


**Supplementary Figure 3: A representative CPG model of respiratory pattern generation does not encode respiratory phase transitions in population activity.**

To illustrate the lack of population coding in respiratory CPG models, we simulated the respiratory CPG model of (49) as described and measured its population firing rate (green). As expected, because phase transitions in CPG models involve an escape or release mechanism, the population firing rate at transitions between respiratory phases was either balanced (PI-E2 transition, red dashed lines) or involved a transition to a new plateau (E2-I and I-PI transitions, blue- and green-dashed lines, respectively).
